# Antagonism and selective modulation of the human glucocorticoid receptor both reduce recruitment of p300/CBP and the Mediator complex

**DOI:** 10.1101/2023.05.15.540854

**Authors:** Laura Van Moortel, Annick Verhee, René Houtman, Diana Melchers, Louis Delhaye, Jonathan Thommis, Kris Gevaert, Sven Eyckerman, Karolien De Bosscher

**Affiliations:** VIB-UGent Center for Medical Biotechnology, 9052 Ghent, Belgium; Department of Biomolecular Medicine, Ghent University, 9052 Ghent, Belgium; Precision Medicine Lab, 5349 AB Oss, The Netherlands

## Abstract

Exogenous glucocorticoids are frequently used to treat inflammatory disorders and as adjuncts for treatment of solid cancers. However, their use is associated with severe side effects and therapy resistance. Novel glucocorticoid receptor (GR) ligands with a patient-validated reduced side effect profile have not yet reached the clinic. GR is a member of the nuclear receptor family of transcription factors and heavily relies on interactions with coregulator proteins for its transcriptional activity. To elucidate the role of the GR interactome in the differential transcriptional activity of GR following treatment with agonists, antagonists, or lead selective GR agonists and modulators (SEGRAMs), we generated comprehensive interactome maps by high-confidence proximity proteomics in lung epithelial carcinoma cells. We found that the GR antagonist RU486 and the SEGRAM Dagrocorat both reduced GR interaction with CREB-binding protein (CBP)/p300 and the Mediator complex when compared to the full GR agonist Dexamethasone. Our data offer new insights into the role of differential coregulator recruitment in shaping ligand-specific GR-mediated transcriptional responses.

**In Brief:** Glucocorticoids are commonly prescribed for the treatment of inflammatory disorders but are associated with severe side effects. Novel glucocorticoid receptor (GR) ligands with strong anti-inflammatory effects but reduced side effects are still sought after. Despite decades-long GR research, there is still an incomplete understanding of the molecular mechanisms driving context-specific GR activity. Using proximity labeling proteomics, we identified CREB-binding protein (CBP), p300 and the Mediator complex as potential crucial GR coregulators driving ligand-induced changes in GR’s transcriptional activity.

**Highlights:**

- Glucocorticoids (GCs), potent anti-inflammatory agents, can elicit side effects
- More selective GCs, causing less side effects, are currently still unavailable
- Lack of fundamental insights on context-specific actions of the GC receptor (GR)
- We mapped ligand-specific GR interactomes using proximity labeling proteomics
- p300/CBP and Mediator undergo ligand-dependent changes in interaction with GR

## Introduction

Synthetic glucocorticoids (GCs) are cheap and extremely potent anti-inflammatory agents that are commonly used to treat inflammatory disorders, severe COVID-19, and as an adjunct in solid cancer treatment (1–4). However, their prolonged use is associated with severe side effects and therapy resistance, which significantly impact patients’ quality of life (5). GCs exert their effect via binding to the glucocorticoid receptor (GR, *NR3C1*), a transcription factor that belongs to the nuclear receptor superfamily. As all nuclear receptors, GR consists of an N-terminal transactivation domain (NTD), a central DNA-binding domain (DBD), and a C-terminal ligand-binding domain (LBD), separated from the DBD by a flexible hinge region (HR) (6, 7).

In the absence of a ligand, GR is predominantly localized in the cytosol, bound by chaperones (8). Upon ligand binding, the receptor dissociates from its chaperone complex and translocates to the nucleus. There, GR has a dual mode-of-action to establish an anti-inflammatory response. On the one hand, GR binds glucocorticoid response elements (GREs) in regulatory DNA elements to mediate transcriptional activation of downstream target genes (9, 10). These target genes elicit a GC-mediated anti-inflammatory response, yet also significantly contribute to metabolic side effects, which are detrimental for the patient. On the other hand, GR can also repress gene transcription, facilitated through its binding to inverted-repeat negative GREs (IR-nGREs) or through functional interaction with pro-inflammatory transcription factors such as nuclear factor kappa B (NF-ĸB) and activator protein (AP)-1. This results in the repression of pro-inflammatory mediators, representing the main mechanism for a GC-mediated anti-inflammatory response (11–13).

GR strongly relies on interactions with coregulatory proteins to exert its transcriptional activity, a process which is facilitated by the formation of GR phase-separated condensates (14). Structurally, GR contains two main coregulator interaction sites, known as activation function (AF)-1 and -2. AF-1 is located in the intrinsically disordered NTD and predominantly responsible for ligand-independent interactions with coregulators and components of the basal transcriptional machinery (15–17). AF-2 is located at the C-terminus of the LBD and facilitates ligand-dependent interactions with coregulators (18, 19). Well-known GR coregulators include the Nuclear Receptor CoActivator (NCoA) and CoRepressor (NCoR) proteins, which assist in transcriptional activation and transcriptional repression, respectively (12, 20). Other complexes contributing to GR-mediated transcriptional regulation include chromatin modifiers and ATPase-dependent chromatin remodelers (21). These alter chromatin organization in such a way that assembly of the RNA polymerase II transcription initiation complex (PIC) is either facilitated (transcriptional activation) or inhibited (transcriptional repression) (22, 23). GR predominantly binds response elements in enhancer regions (24) and therefore relies on interaction with the Mediator complex to transfer regulatory signals from enhancer regions to the PIC at core promoters (25). GR actions are further fine-tuned by coregulator post-translational modifications (PTMs) (26).

The significant side effect burden of GCs has led to the search for more selective GR ligands, so-called selective GR activators and modulators (SEGRAMs). These ligands were designed to reduce the expression of metabolic genes via reduced GR activity on GREs while maintaining its ability to interfere with NF-ĸB and AP-1 activity (27). However, despite extensive research, none of the lead SEGRAMs reached the clinic (reviewed in (28)). In a recent study, we characterized a panel of well-known SEGRAMs and found that most of them behaved as (weak) agonists, with reduced GR transcriptional activity (29). We also found that both SEGRAMs and GR antagonists reduced the levels of GR Ser211 phosphorylation, a PTM which has been reported to facilitate the AF-1-mediated interaction with coregulator proteins (30). This observation led us to question whether and how such ligands would affect the GR interactome compared to full agonist and antagonist activities, and whether observed changed might explain the reduced transcriptional activity of GR.

While the global interactome of agonist-bound GR in human embryonic kidney (Hek293T) cells and murine bone marrow-derived macrophages (BMDMs) has been resolved (31–33), studies with SEGRAMs have been limited to the binary mapping of a handful of GR-protein interactions (34–36). In our current study, we used proximity-dependent biotin identification (BioID) with an improved control system in human epithelial carcinoma (A549) cells to map the global interactome of GR bound by the full agonist Dexamethasone (Dex), the antagonist RU486 or a well-studied SEGRAM, Dagr (36, 37). We found that RU486 and Dagr reduced GR interaction with NCoA2 and essential coregulators for enhancer-based transcriptional activation, such as CREB-binding protein (CBP), p300 and the Mediator complex (25, 38, 39).

## Experimental procedures

### Materials

Dexamethasone (Dex, D4902) was purchased from Sigma-Aldrich. RU486 and Dagr were kind gifts. All compounds were dissolved in DMSO, aliquoted and stored at -20 °C at a stock concentration of 10 mM. Dex and Dagr were used at a final concentration (f.c.) of 1 µM, RU486 was used at a f.c. of 0.1 µM. Doxycycline hyclate (Sigma, D9891) was dissolved in MQ, aliquoted and stored at -20 °C in the dark. Biotin (Sigma, B4639) was dissolved in 60 mM NaOH and stored at -20 °C in the dark. Benzonase nuclease was purchased from Sigma (E1014-25KU) and streptavidine sepharose high performance beads (5 mL) was purchased from VWR (GE Healthcare 17-5113-01). OMIX C18 tips (100 µL) were ordered from Agilent (A57 003 100). Mouse anti-β-actin antibody (A1978) was purchased from Sigma Aldrich. Anti-GST Alexa488 antibody (A11131), recombinant GR-LBD-GST (A15668) and Nuclear Receptor Buffer F (PV4547) were all from Invitrogen.

### Cell culture

All A549-derived cell lines were cultured in Dulbecco’s Modified Eagle Medium (DMEM) supplemented with 10% fetal bovine serum (FBS) and 5 µg/mL blasticidine.

### Stable cell line generation

The generation of the A549_1128 cell lines expressing inducible V5-TurboID-T2A-GR and V5-Turbo-mutT2A-GR is described in (40). In brief, 20,000 A549_1128 cells were cultured in a 24-well plate. 48 h later, they were transfected with 100 ng pCMV-hyPBAse (coding for hyperactive Piggybac transposase) and 400 ng pPB-TRE_V5-TurboID-T2A-GR or pPB-TRE_V5-TurboID-mutT2A-GR using lipofectamine LTX (ThermoFisher Scientific). Two days after transfection, the cells were transferred to a T25 falcon, and five days after transfection, 5 µg/mL blasticidine was added to the culture medium. Cells were expanded in selection medium and the cell pools were used for further experiments.

### TurboID experiments

For every condition, three 15-cm culture dishes with 2.7 x 10^6^ A549 V5-TurboID-T2A-GR or A549 V5-TurboID-mutT2A-GR cells were seeded in DMEM supplemented with 10% FBS and 10 µg/mL gentamicin (4 replicates per condition). 24 h later, 150 pg/µL doxycycline was added to the A549 V5-TurboID-T2A-GR cells, and 20 pg/µL was added to the A549 V5-TurboID-mutT2A-GR cells. 48 h after seeding, 50 µM biotin was added to each plate, in combination with either solvent, Dex, RU486 or Dagr. Cells were incubated for 2 h, after which they were put on ice and washed with DPBS followed by detachment in 1 mL DPBS using a cell scraper (Corning). Cells from the same condition (3 plates) were pooled in a 15 mL tube and pelleted for 5 min at 500g (4 °C). Next, they were washed in 6 mL DPBS before storing the cell pellets at - 20 °C. The next day, cell pellets were lysed on a rotator for 1 h (4 °C) in 2.5 mL ice-cold RIPA lysis buffer (50 mM Tris-HCl pH 7.5, 150 mM NaCl, 1% NP-40, 2 mM EDTA pH 8, 0.1% SDS, 0.5% sodium deoxycholate, EDTA-free cOmplete protease inhibitor cocktail) supplemented with 1 µL benzonase nuclease. Next, samples were sonicated using a stick sonicator (five 6-second bursts, amplitude 30) before removing cellular debris via centrifugation (15 min at 29,000g, 4°C). Supernatant was transferred to a new tube, protein concentrations were determined using the Pierce™ BCA Protein Assay (ThermoFisher) and equalized across samples. Streptavidin sepharose beads were pre-washed three times in lysis buffer without DOC or protease inhibitor. Next, 90 µL beads was incubated with 1800 µg lysate on a rotator for 3 h (4 °C), followed by three washing step in RIPA buffer, two washing steps with 50 mM ammonium bicarbonate pH 8 and one washing step in 20 mM Tris-HCl pH 8, 2 mM CaCl_2_. Beads were resuspended in 100 µL 20 mM Tris-HCl pH 8 before overnight digestion with 1 µg trypsin in a Thermoshaker (37 °C, 1200 rpm). The next day, digested proteins were transferred to a new LoBind Eppendorf before further digesting further with 0.5 µg trypsin for 3 h (37 °C, 750 rpm). Finally, samples were acidified to 2% formic acid (FA) before performing peptide clean-up using Agilent OMIX C18 100 µL tips (A57 0003 100) with pre-wash using 80% acetonitrile (ACN), 0.1% trifluoroacetic acid (TFA); wash steps with 20% ACN, 0.1% TFA and elution using 60% ACN, 0.1% TFA. Samples were dried in a SpeedVacTM vacuum concentrator and stored at -20 °C until further use.

### LC-MS/MS

Peptides were re-dissolved in 20 µL loading solvent A (98% ACN, 0.1% TFA) of which 3.5 µL was injected for LC-MS/MS analysis on an Ultimate 3000 RSLCnano system in-line connected to a Q Exactive HF mass spectrometer (Thermo Fisher Scientific). Trapping was performed at 20 μL/min for 2 min in loading solvent A on a 5 mm trapping column (Thermo scientific, 300 μm internal diameter (I.D.), 5 μm beads). The peptides were separated on a 250 mm Aurora Ultimate, 1.7 µm C18, 75 µm inner diameter (IonOpticks) kept at a constant temperature of 45 °C. Peptides were eluted by a non-linear gradient starting at 1% MS solvent B (80% ACN, 0.1% FA) reaching 33% MS solvent B in 60 min, 55% MS solvent B in 75 min, 70% MS solvent B in 90 minutes followed by a 10-minute wash at 70% MS solvent B and re-equilibration with MS solvent A (0.1% FA). QCloud has been used to control instrument longitudinal performance during the project (41, 42). The mass spectrometer was operated in data-dependent mode, automatically switching between MS and MS/MS acquisition for the 12 most abundant ion peaks per MS spectrum. Full-scan MS spectra (375-1500 m/z) were acquired at a resolution of 60,000 in the Orbitrap analyzer after accumulation to a target value of 3,000,000. The 12 most intense ions above a threshold value of 15,000 were isolated with a width of 1.5 m/z for fragmentation at a normalized collision energy of 30% after filling the trap at a target value of 100,000 for maximum 80 ms. MS/MS spectra (200-2000 m/z) were acquired at a resolution of 15,000 in the Orbitrap analyzer.

### Data analysis

Data searching was done with the MaxQuant software (v1.6.11.0) using the Andromeda search engine with default search settings, including a false-discovery rate (FDR) of 1% on both the peptide and protein level. Spectra were searched against the human SwissProt proteome database (version of May 2021, 20386 entries). The FASTA sequence of V5-TurboID was added to the search database. The mass tolerance for precursor and fragment ions was set to 20 ppm and 4.5 ppm, respectively. Enzyme specificity was set as C-terminal to arginine and lysine (trypsin), also when followed by a Pro residue, with a maximum of two missed cleavages. Variable modifications were set to oxidation of methionine residues and acetylation of the protein N-terminus; no fixed modifications were set. A minimum of one razor or unique peptide was required for identification. Matching between runs was enabled with a 20-min alignment window and a matching time window of 0.7 min. Proteins were quantified by the MaxLFQ algorithm integrated in the software, with the fastLFQ switched off and a minimum ratio count of two unique or razor peptides. An overview of all protein and peptide identifications as generated by MaxQuant can be found in **Table S1** (Sheet 1 contains the Peptides.txt output file, Sheet 2 contains the ProteinGroups.txt output file). Further data analysis was performed with the Perseus software (v.1.6.15.0) using the ProteinGroups table from the MaxQuant search output. Potential contaminants and proteins identified in a reverse database were removed, as well as proteins that were only identified by site. The protein LFQ intensities were log_2_-transformed to obtain a normal distribution (**Figure S1**) and the data were analyzed further for each ligand separately. For each comparison, proteins with less than three identifications in at least one group were removed. Missing values were replaced by imputation from a normal distribution (width 0.3, downshift 1.8), and significantly enriched or depleted proteins were identified via two-sample t-tests with a permutation-based FDR (1000 randomizations) of 1% or 5% and an S_0_ of 0.58 for truncation. Volcano and dotplots were visualized using the ggplot2 package in RStudio. For gene ontology functional enrichment of each condition, the UniProt identifiers of all proteins and their difference in log_2_(LFQ intensities) were imported in STRING. The Uniprot identifiers of all significantly enriched proteins (5% FDR, S_0_ = 0.58) and their difference in log_2_(LFQ intensity) were used to create physical networks in STRING, which were processed further using Cytoscape.

### Immunoblot analysis

18 µg input or unbound fraction and 8 µL of the IP fraction of each TurboID sample was loaded on a 10% polyacrylamide gel. Following SDS-PAGE, proteins were transferred to 0.45-µm nitrocellulose membrane (GE Healthcare) for 2 h at 100V. Membranes were blocked using StartingBlock TBS blocking buffer (Thermo Scientific) mixed 1:1 with TBS-T (50 mM Tris-HCl pH 7.5, 150 mM NaCl supplemented with 0.1% Tween20) and subsequently incubated with antibodies against β-actin (1:20,000) and biotin (1:10,000).

### Nuclear receptor Activity Profiling (NAPing)

GR-coregulator binding, as parameter for receptor conformation and activity status, was profiled using the NAPing platform (PML, Oss, The Netherlands). In short, a reaction mix of GST-tagged GR LBD (final concentration of 10 nM), fluorescently labeled GST-antibody (final concentration of 25 nM) and Nuclear Receptor Buffer F (A15668, A11131, PV4547, Thermo Fisher Scientific) supplemented with 1 μM compound or solvent only (DMSO, 2% final concentration) was incubated with a set of 101 immobilized 20-mer peptides, each representing a unique coregulator-derived NR-binding motif. After 60 min, unbound receptor and antibody were removed by washing and, subsequently, NR-coregulator binding (fluorescence) was quantified by in-house dedicated software R-scripts.

Each condition was analyzed using three technical replicates, resulting in GR-coregulator binding, mean ± S.E.M.. Significant changes in GR-coregulator interaction across conditions was evaluated using one-way ANOVA, followed by Sidak’s multiple comparisons with correction for multiple testing.

### Experimental Design and Statistical Rationale

The TurboID experiments were conducted with four biological replicates for each condition (32 samples in total), as we expected limited biological variation between replicates in the immortalized cell lines. We included a control cell line expressing sole TurboID in order to distinguish true from aspecific interactors. A DMSO control condition was included to correct for solvent effects (four replicates). Each ligand (or solvent) treatment was conducted on control and target cell lines to correct for ligand-induced changes in protein levels. For the same reason, two-sided t-tests (1% FDR, S_0_ = 0.58) were performed on control versus target cell lines for each ligand treatment separately, rather than directly comparing the ligand-specific interactomes generated in the target cell line. The histograms in **Figure S1** confirm a normal distribution of the data. The order of the samples was randomized during the sample preparation; the LC-MS/MS runs were performed in randomized blocks.

*In vitro* NAPing experiments were performed with three technical replicates for each condition (12 samples in total), as we expected minimal technical variation between replicates. A DMSO control condition was included to correct for solvent effects (3 replicates). Changes in GR-peptide binding between different ligands were evaluated via one-way ANOVA, followed by Sidak’s multiple comparisons.

## Results

### TurboID proximity labeling combined with a T2A split/link design yields high-confidence GR interactomes

To evaluate the role of the interactome in GR’s reduced transcriptional activity with SEGRAMs and antagonists, we mapped the GR interactome following 2 h stimulation with Dex, RU486 or Dagr using proximity-dependent biotin identification (BioID). To ensure optimal compatibility with 2 h biotin labeling times, we opted to use TurboID, a more active variant of the original BirA* biotin ligase (43). Using the transposon-based PiggyBac system (44), we generated two A549 cell lines with stably integrated TurboID-GR under a doxycycline-inducible promoter. To distinguish true from aspecific interactors, we made use of the T2A split/link design (**Figure 1**) (45). In the control cell line, TurboID and GR were separated by a T2A tag, leading to ribosomal skipping and the expression of TurboID and GR as two individual proteins. In the target cell line, the T2A tag was mutated and inactivated (mutT2A), resulting in the expression of a TurboID-mutT2A-GR fusion protein. Doxycyclin concentrations were optimized to match bait protein expression between both set-ups at near-endogenous levels. The generation and characterization of these cell lines has been described in detail in (40). 24 h after induction of bait expression, we treated both cell lines with ligand and 50 µM biotin for 2 h and performed streptavidin enrichment to extract biotinylated proteins. Western blot controls confirmed comparable biotinylation profiles across all conditions (**Fig. S2**). Next, LC-MS/MS was used to identify and quantify enriched proteins. Using protein label-free quantification, comparing protein intensities between target and control cell lines, we identified the GR interactome for each of the ligand treatments (**Figure 2**). High quality data sets were obtained, with GR (*NR3C1*) significantly enriched in all four conditions (**Figure 2**). At a 1% FDR level, we found 12 significantly enriched proteins in the solvent condition, and identified 125, 88 and 118 GR interaction partners with Dex, RU486 and Dagr, respectively (**Figure 2B-D**, **Fig. S3-S6**, **Table S2**). 77 of these interactors were shared by all ligands and 52 of all identified proteins have already been identified as GR interactors according to BioGrid (**Table S2**) (46). Overall, our data confirmed that a combination of TurboID with the T2A split/link design yielded high-confidence GR interactomes following 2 h biotin labeling.

**Figure 1.**
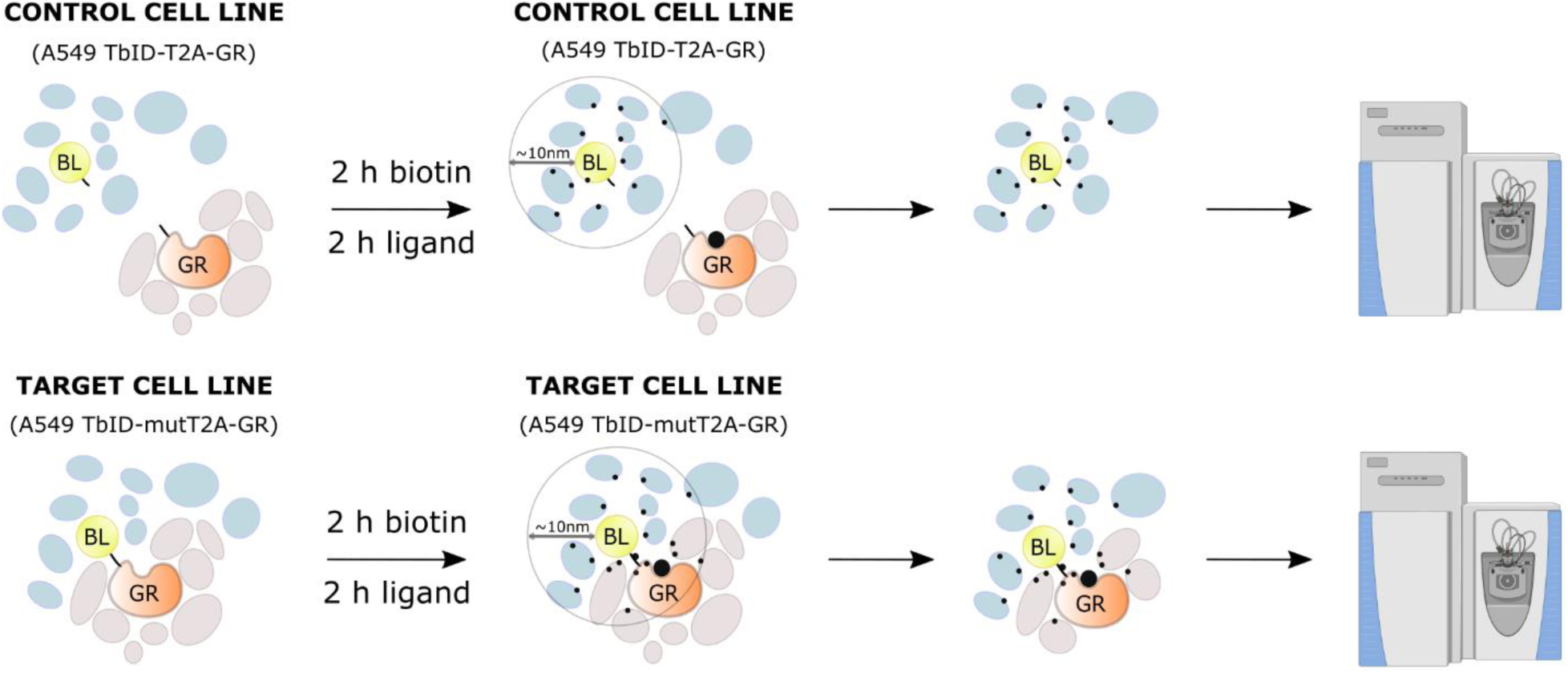
Overview of the TurboID T2A split/link design. In the control cell line, TurboID and GR are separated by a T2A tag, leading to their expression as two separate proteins. In the target cell line, TurboID-mutTA-GR is expressed as a single protein due to T2A inactivation. Direct comparison of proteins identified in the target versus control cell line allows distinction between aspecific interactors and GR interactome components. BL, biotin ligase; GR, glucocorticoid receptor; TbID, TurboID.

**Figure 2.**
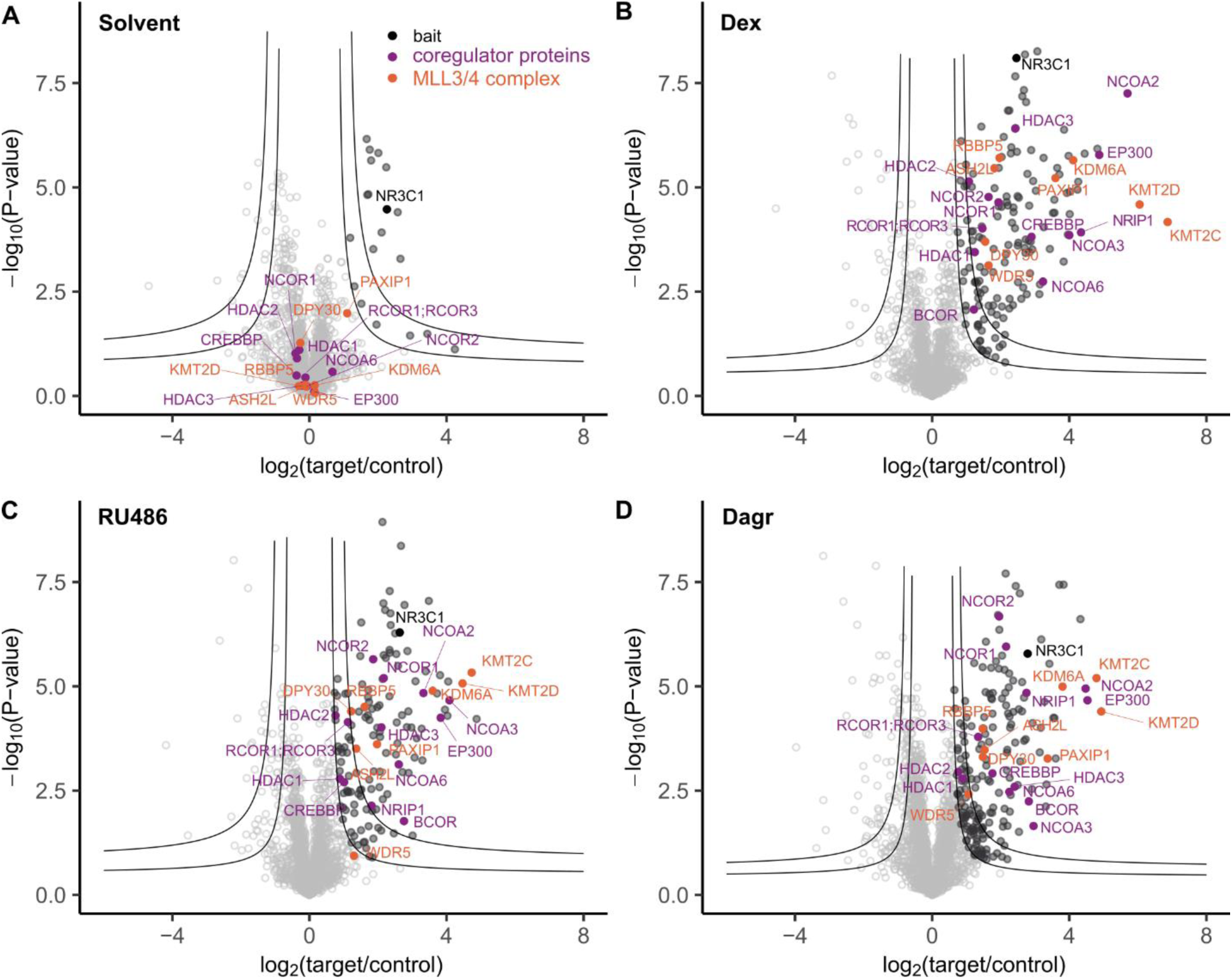
Volcano plots representing changes in log_2_(LFQ intensity) (x-axis) and -log_10_(P values) (y-axis) between target and control cell lines for each condition. The two cut-offs represent S_0_ = 0.58 in combination with 5% or 1% FDR. Full circles indicate significantly enriched proteins at the 5% FDR level. Bait is annotated in black, known GR interactors in purple and components of the MLL3/4 complex in orange.

### Agonist-induced GR recruits the Mediator complex and chromatin remodelers and modifiers

We found many well-known direct GR interactors in the Dex condition, including not only the coactivators NCoA2 and NCoA3, but also the corepressors NCoR1 and NCoR2 (**Figure 3**, **Table S2**). In addition, we found several more recently identified GR interactors that were also found in previously published GR interactome studies in human embryonic kidney cells, such as Bcl-6 corepressor (BCOR), REST corepressor (RCOR)1 and interferon regulatory factor 2 binding protein 2 (IRF2BP2) (31). Next, we used all significantly enriched proteins in the Dex condition at a 1% FDR as input for gProfiler functional enrichment analysis (https://biit.cs.ut.ee/gprofiler/gost). We identified 134 enriched protein complexes in the integrated CORUM database (47), which showed substantial overlap in composition (**Table S4**). The Mediator complex was most significantly enriched (**Figure 3**, **Table S4**). We also identified several other protein complexes associated with transcriptional regulation, including the ATPase-dependent chromatin remodeler Switch/Sucrose non-fermentable (SWI/SNF) and the MLL3/4 histone N-lysine methyltransferase (KMT) complexes (**Figure 2**, **Table S4**). KMT2C and KMT2D, the catalytic subunits of the MLL3/4 complexes, were even the two most enriched proteins in the Dex condition (**Figure 2B**, **Table S2**). Interestingly, many components of the (expected nuclear) SWI/SNF complex were also significantly enriched in the solvent condition (**Table S2, S3**). We also found several complexes with histone acetyltransferase (HAT) activity in the agonist-bound GR interactome, which are also involved in chromatin reorganization. These HATs include the Nucleosome acetyltransferase of H4 (NuA4) and Spt-Ada-GCN5 (SAGA) complexes (**Figure 3**, **Table S4**). Reassuringly, we identified p300 and CBP, two other well-known GR interactors with HAT activity (30, 48, 49). Finally, we identified several components of the Nucleosome Remodeling and Deacetylase (NuRD) complex, which possess ATP-dependent chromatin remodeling as well as histone deacetylase activities (50).

**Figure 3.**
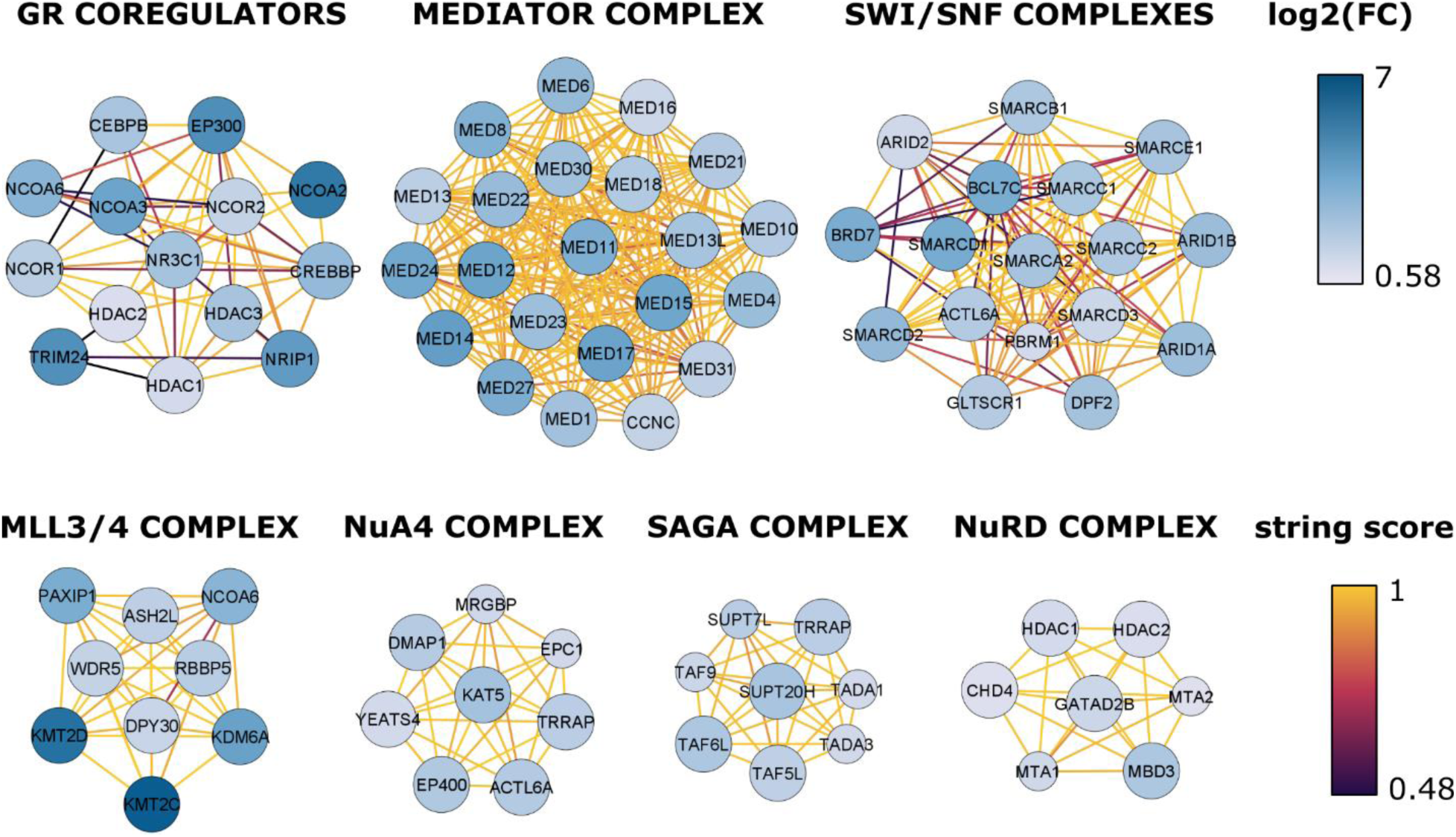
Identified protein complexes associated with Dex-induced GR. Small circles represent significant interactors at a 5% FDR, larger circles represent significant interaction partners at a 1% FDR. The blue color scale indicates the log_2_(fold change) between target and control cell line. The purple-yellow color scale corresponds to the confidence of the interaction according to the STRING database.

### Antagonists and partial agonists cause reduced GR recruitment of NCoA2, p300/CBP and the Mediator complex

Next, we compared the relative enrichment of the protein complexes shown in **Figure 3** between solvent and the three ligands at a 1% FDR. The majority of the typical GR coregulator proteins, including NCoA3, NCoA6, NCoR1 and NCoR2, showed comparable enrichments between the different conditions (**Figure 4** **and S7**). However, we did find a pronounced increase in NCoA2 recruitment with Dex, compared to RU486 and Dagr, and reduced interaction with the recently identified GR corepressor BCOR. In addition, we also found increased GR interaction with the corepressor protein Nuclear Receptor Interacting Protein (NRIP)1 in the Dex condition (**Figure 4** **and S7**).

**Figure 4.**
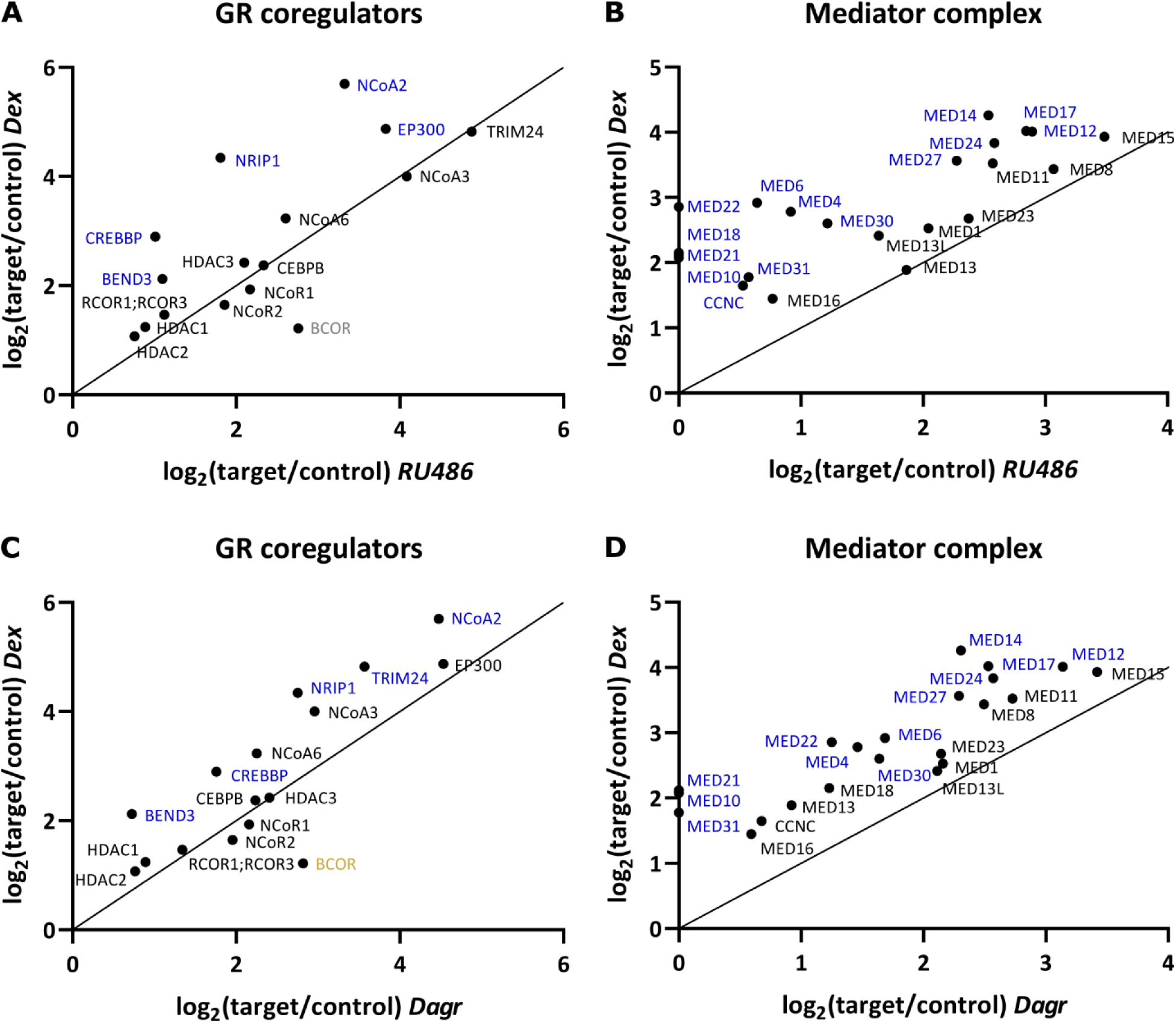
Changes in enrichment of GR coregulators and Mediator subunits between Dex, RU486 and Dagr. **(A)** Comparison of GR coregulator enrichments between Dex and RU486. **(B)** Comparison of Mediator subunit enrichments between Dex and RU486. **(C)** Comparison of GR coregulator enrichments between Dex and Dagr. **(D)** Comparison of Mediator subunit enrichments between Dex and Dagr. Proteins indicated in blue are at least two-fold more enriched in the Dex condition, proteins in grey are at least two-fold more enriched in the RU486 condition, proteins in yellow are at least two-fold more enriched in the Dagr condition.

We observed a remarkable change in the GR interaction with the Mediator complex between the different ligands, as indicated by the changes in Mediator complex enrichment in gProfiler analyses (**Figure 4B, 4D, Tables S4-S6**). Several Mediator subunits that were identified with Dex, were not found in the dataset with RU486 (e.g., MED10, MED18, MED21 and MED22) or were not significantly enriched at a 1% FDR (e.g., CCNC, MED4, MED6, MED16, MED30 and MED31). Other subunits were still significantly enriched but with a markedly lower fold change than in the Dex condition (e.g., MED14, MED17 and MED24). Mediator recruitment was also decreased with Dagr compared to Dex, although the difference was slightly less pronounced than with RU486 (**Figure 4D**).

We found only mild changes in the fold enrichment of most chromatin remodeling and modifying complexes between the different ligands, even though some proteins were no longer found as significant in the RU486 and Dagr conditions (**Figure S7C-G**). However, we observed a reduced recruitment of CBP (CREBBP) after RU486 treatment compared to Dex (**Figure 4A, 4C, S7A**). Although less pronounced, we made a similar observation for p300.

We also identified many other transcription factors (TFs) which generally changed little in enrichment between the different conditions (**Fig. S8**). Exceptions are TCF12 and ZNF592, which were clearly more enriched in the Dex condition (**Fig S8**).

### Nuclear receptor Activity Profiling on GR LBD confirms ligand-dependent differential GR interactions with Mediator and nuclear receptor coregulators

To validate the results from the BioID experiments, we performed Nuclear receptor Activity Profiling (NAPing). In this assay, recombinant GR-LBD-GST was combined with different ligands and subsequently allowed to bind 101 peptides of well-known NR coregulators, immobilized on a solid support. Given the *in vitro* context of NAPing, this assay allows identification of coregulators that have the ability to bind GR directly, without taking competition with other coregulators or cell-specific changes in coregulator expression levels into account. In addition, by only using the GR LBD, we could also determine which coregulators are able to interact with the AF-2 domain of GR.

The resulting coregulator binding profiles confirmed the ligand-dependent changes in GR interaction with NCoA2 and NRIP1, but also showed marked changes in other interaction profiles. For instance, NCoA1, Peroxisome proliferator-activated receptor gamma coactivator 1-alpha (P(R)GC1α) and nuclear receptor subfamily 0 group B member 2 (NR0B2) were among the most predominant binding peaks in the NAPing profile with Dex (**Figure 5**) but were not identified in any of the BioID datasets. Interestingly, we also found that RU486 and Dagr reduced the interaction GR LBD with NCoA3 compared to Dex, even though such ligand-dependent changes were not observed in our BioID datasets (**Figure 3, 5**). Concerning corepressors and also in contrast to our BioID data, RU486-bound GR LBD showed a more pronounced interaction with NCoR1 and NCoR2 corepressor peptides compared to Dex. Intriguingly, GR LBD barely interacted with any of the NCoR1 and NCoR2 peptides in the Dex and Dagr NAPing conditions, although both proteins were significantly enriched in the BioID data. We also found very little GR LBD interaction with KMT2C, KMT2D, CBP or p300, even though the former two were among the most enriched proteins in the BioID dataset with all three ligands (**Figure 2, 5**).

**Figure 5.**
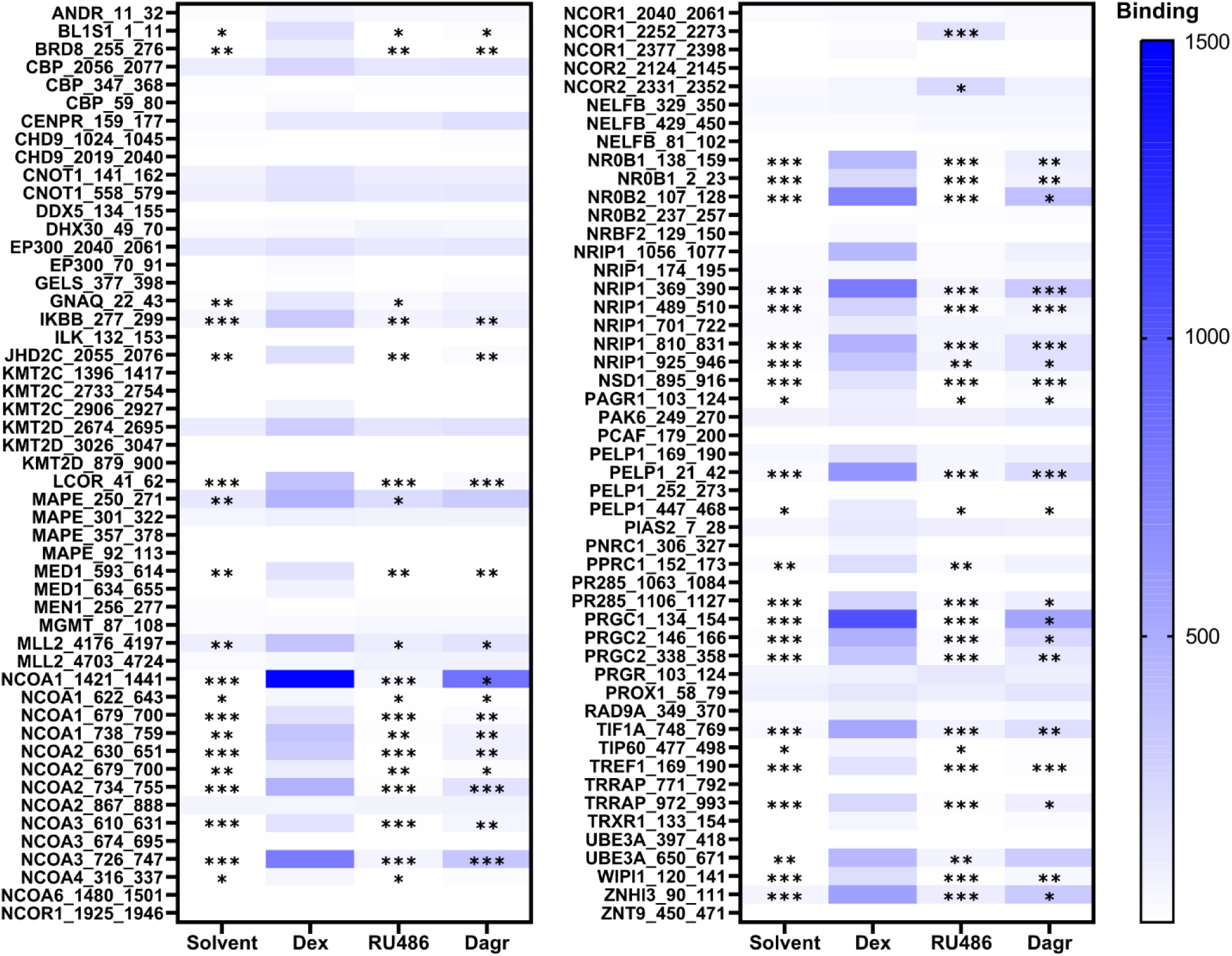
Relative quantification of GR interaction with coregulator peptides using Nuclear receptor Activity Profiling (NAPing) on recombinant GR LBD. The blue color scale indicates the mean GR LBD binding strength as measured by fluorescence intensity (three replicates). Statistically significant changes versus Dexamethasone (Dex) were evaluated via ordinary one-way ANOVA with Sidak’s post-hoc testing (* P < 0.05; ** P < 0.01; *** P < 0.001).

We retrieved only mild, albeit ligand-dependent interaction of GR with MED1 in the NAPing data (**Figure 5**). This was in line with the BioID datasets, where this mediator subunit also showed weaker interaction with GR than other mediator subunits, such as MED14 (**Figure 3B, S7**).

## Discussion

In this study, we mapped the GR interactome following 2 h treatment with solvent, a GR agonist, an antagonist or a SEGRAM. Our previous work has revealed that SEGRAMs and antagonists reduced GR Ser211 phosphorylation and transcriptional activity while supporting a similar GR nuclear translocation profile as full agonists (29). Using BioID, we aimed to elucidate whether changes in the GR interactome could also help explain this reduction in GR’s transcriptional activity, as we hypothesized that reduced GR Ser211 phosphorylation would lead to reduced AF-1-mediated coregulator interactions (30, 51). We used TurboID instead of the less active BirA* biotin ligase to reduce labeling times from 24 h to 2 h. We confirmed that 2 h labeling with TurboID was sufficient to identify many of the well-known GR interactors which were also found in earlier GR proximity mappings using BirA* and 24 h labeling (31, 32). In addition, we found that the GR antagonist RU486 and the SEGRAM Dagr reduced GR interactions with several coregulator proteins, including NCoA2, CBP/p300 and the Mediator complex (**Figure 4, Fig. S7**). To our knowledge, this is the first study reporting global interactome changes that may be in line with a reduced transcriptional activity of GR with SEGRAMs.

When evaluating the global GR interactome across the different ligands using BioID, we found that KMT2C and KMT2D, the catalytic subunits of the histone lysine methyltransferase MLL3/4 complex, were among the most enriched proteins with all three ligands (**Figure 2B-D**). Interestingly, this was not reflected by the NAPing data (**Figure 5**), indicating that the interaction between GR and KMT2C/D might either be indirect, or mediated via the AF-1 in the GR NTD. KMT2C and KMT2D belong to the KMT2 histone lysine methyltransferase subfamily, which comprises six proteins that catalyze the methylation of histone 3 lysine 4 (H3K4), a hallmark of transcriptionally active promoters and enhancers (52). Interestingly, the enrichment of KMT2C and KMT2D but none of the four other members of the KMT2 family is in line with the reported predominant GR binding to enhancers (24, 53), as KMT2C and KMT2D predominantly facilitate H3K4 methylation in enhancer regions (54). We also identified KMD6A, NCOA6 and PAXIP1 (**Figure 3, S7**), which are components of the MLL3/4 complex but not of the complexes comprising other KMT2 subfamily members (54). Our finding is in contrast to another recent GR interactome dataset in BMDMs, which was generated using ChIP-MS and where KMT2F and KMT2G were the most predominant KMT2 family members (33). This discrepancy might be explained by either cell type specific differences in coregulator content or ratios, or alternatively, by the absence of an inflammatory context in our experiments. An inflamed cell status was shown to result in altered GR-chromatin binding profiles (55) that may also lead to differences in the overall recruitment of particular GR coregulators.

We identified several NCoA proteins that are known to bind GR directly via LXXLL motifs (**Figure 3, 4)** (56–59). Remarkably, we did not identify NCoA1 in our BioID dataset, although this protein was identified in several other GR interactome studies (31, 32, 60) and was one of the most prominent GR interactors in our NAPing dataset. The absence of this coregulator in our BioID datasets could potentially be explained by its low protein expression levels in A549 cells under high-glucose conditions (61). PGC1α is another well-known GR coregulator that was not identified in our BioID data. However, this protein was also not identified in GR interactome datasets in human embryonic kidney cells or BMDMs (31–33), which may correlate with a most predominant expression in liver, muscle and brown adipose tissue (62). NCoA2 was the only NCoA protein showing differential interaction with GR following antagonist or SEGRAM binding, both in the BioID and NAPing datasets (**Figures 4, 5 and S7**). This differential interaction is in line with existing structural data on the GR LBD with different ligands, which indicate that GR antagonists and SEGRAMs alter the conformation of the LBD and AF-2 in such a way that NCoA2 binding is sterically hindered (63, 64). In addition, reduced GR-NCoA2 interaction with Dagr and RU486 has also been shown in a study using mammalian two-hybrid assays (36). The differential GR interaction with NCoA2, together with the reported reduction in the phosphorylation of Ser211 (29), could help to explain our observed differences in GR interaction with the HATs p300 and CBP across the different ligands (**Figure 4A, 4C and S7**). More specifically, p300 and CBP have both been shown to interact directly with several NCoA proteins (57, 65, 66), but they can also interact with the AF-1 region of nuclear receptors (67, 68). Indirect or AF-1-mediated interaction between GR and p300/CBP is in line with the absence of p300 and CBP interaction with the GR LBD in our NAPing data (**Figure 5**). Moreover, CBP interaction with GR increases upon Ser211 phosphorylation (30). p300 and CREB-binding protein (CBP) catalyze the acetylation of H3K27, a hallmark of active enhancers and required for long-range interactions with the basal transcriptional machinery (38, 39, 69, 70). Furthermore, the induction of H3K27ac by p300 and CBP is also essential for the recruitment of RNA polymerase II and subsequent transcriptional activation (13, 71). Therefore, the reduced GR interaction with p300 and especially CBP following treatment with RU486 and Dagr (**Figure 4A, 4C**), could help to explain why GR shows reduced transcriptional activity with these ligands.

Another component that is essential for long-range interactions between enhancers and the basal transcriptional machinery is Mediator (25). We found reduced GR interaction with many components of this complex in the antagonist and SEGRAM conditions. This effect appeared to be most pronounced for MED14, the Mediator subunit which serves as the backbone for this complex (**Figure 4, S7**) (25). MED14 was shown to interact directly with GR via its AF-1 (72, 73), which is also in line with our hypothesis that reduced GR Ser211 phosphorylation leads to reduced AF-1-mediated coregulator interactions. Over the years, several studies have indicated that GR’s interaction with another Mediator subunit, MED1, is mediated via AF-2 and relies on the coregulator’s two LXXLL motifs (74). This was confirmed by our NAPing data, which indeed revealed interaction of Dex-bound GR LBD with MED1 (**Figure 5**). Although this interaction was weak in all conditions, we were still able to find significant, ligand-dependent changes, in line with our BioID data.

Besides diminished transcriptional activation of GR, we also aimed to assess whether our interactome data could provide a plausible explanation for the reduced transcriptional repression capacity of GR when bound by SEGRAMs and antagonists, indicated by reduced repression of inflammatory mediators and reduced IR-nGRE-mediated downregulation of GR levels (29). Two studies by Hua and colleagues have shown that NCoR1, NCoR2 and HDAC3 are all essential for GR-driven inhibition of NF-ĸB and AP-1, and for IR-nGRE mediated transcriptional repression by GR in mouse embryonic fibroblasts (12, 75). However, surprisingly, we found no changes in GR interaction with either of these three proteins between the different ligand treatments in A549 cells (**Figure 4A, 4C**). This indicates that the role of NCoR1, NCoR2 and HDAC3 in GR-mediated transcriptional repression might be cell type-specific, or that these coregulator proteins are preferentially recruited to chromatin regions that are only GR-bound under inflammatory conditions (55). However, earlier interactome studies in human embryonic kidney cells also showed increased interaction between NCoR1 and Dex-associated GR compared to RU486-associated GR (31) in absence of an inflammatory context, arguing against the latter hypothesis and indicating that cell type might have a more considerable influence than an inflammatory context. Even more puzzling, we did find an enhanced interaction of antagonist-bound GR LBD with NCoR1 and NCoR2 in the NAPing data (**Figure 5**). This finding is in line with data from GR LBD crystallization studies, which indicate that GR LBD bound by an antagonist favors interaction with NCoRs over interaction with NCoA proteins (63). The discrepancy between the BioID and NAPing outcomes for the corepressors indicates that GR’s interaction with NCoR proteins might not only be mediated by AF-2, but also by AF-1. This is supported by the studies of Hua and colleagues, wherein ChIP-qPCR on IR-nGRE and tethering GR targets revealed that truncation of the GR NTD abrogated recruitment of NCoR1 and NCoR2 to GR-bound chromatin regions (12, 75).

Future studies involving more GR ligands with diverse activity profiles should confirm whether the differential GR interaction with NCoA2, p300/CBP and Mediator is indeed linked to GR transcriptional activity. Furthermore, interactome studies using a GR Ser211 mutant could yield further insights into the role of defective Ser211 phosphorylation in GR coregulator interactions. In addition, follow-up studies should also be expanded to other cell types, as we expect that ligand-dependent changes in GR coregulator expression profiles will also differ between cell types. Finally, ChIP-or CUT&RUN-sequencing studies could help elucidate if reduced chromatin binding of GR also contributes to its reduced transcriptional activity with antagonists and SEGRAMs. However, since the SWI/SNF complex and most chromatin-modifying complexes show no ligand-dependent differences in association with GR, we predict that GR binding profiles may be comparable.

In conclusion, our data support that a differential coregulator protein recruitment profile may play a role in shaping ligand-specific GR-mediated transcriptional responses. Herewith we contribute to the quest for a more refined GR coregulator profile that should ultimately match a better-delineated GR transcriptional profile.

## Data availability

The mass spectrometry proteomics data have been deposited to the ProteomeXchange Consortium via the PRIDE partner repository (76) with the dataset identifier PXD042014 and 10.6019/PDX042014. The NAPing data are to be shared upon request to Prof. Karolien De Bosscher.

## Conflicts of Interests

none

## Acknowledgments

We thank Francis Impens, Simon Devos and Sara Dufour from the VIB Proteomics Core for the LC-MS/MS analysis. We thank Dorien Clarisse and Lisa Koorneef for their assistance with the BioID sample preparation. We thank George Moschonas for his help with the visualization of the protein networks.

## Supplemental data

This article contains Supplemental data.

## Funding

This work was funded by a Research Foundation Flanders (FWO) grant to Sven Eyckerman (G042918N). Louis Delhaye is supported by a UGent doctoral-assistant mandate (BOF22/PDO/024). Laura Van Moortel was supported by a strategic basic research fellowship of the Research Foundation Flanders, grant number 1S14720N. The funding sources had no role in the study design, data collection, analysis and interpretation, nor in the writing process.

## Notes

### Competing Interest Statement

The authors have declared no competing interest.

