## Supplementary figures and tables for "Antagonism and selective modulation of the human glucocorticoid receptor both reduce recruitment of p300/CBP and the Mediator complex"

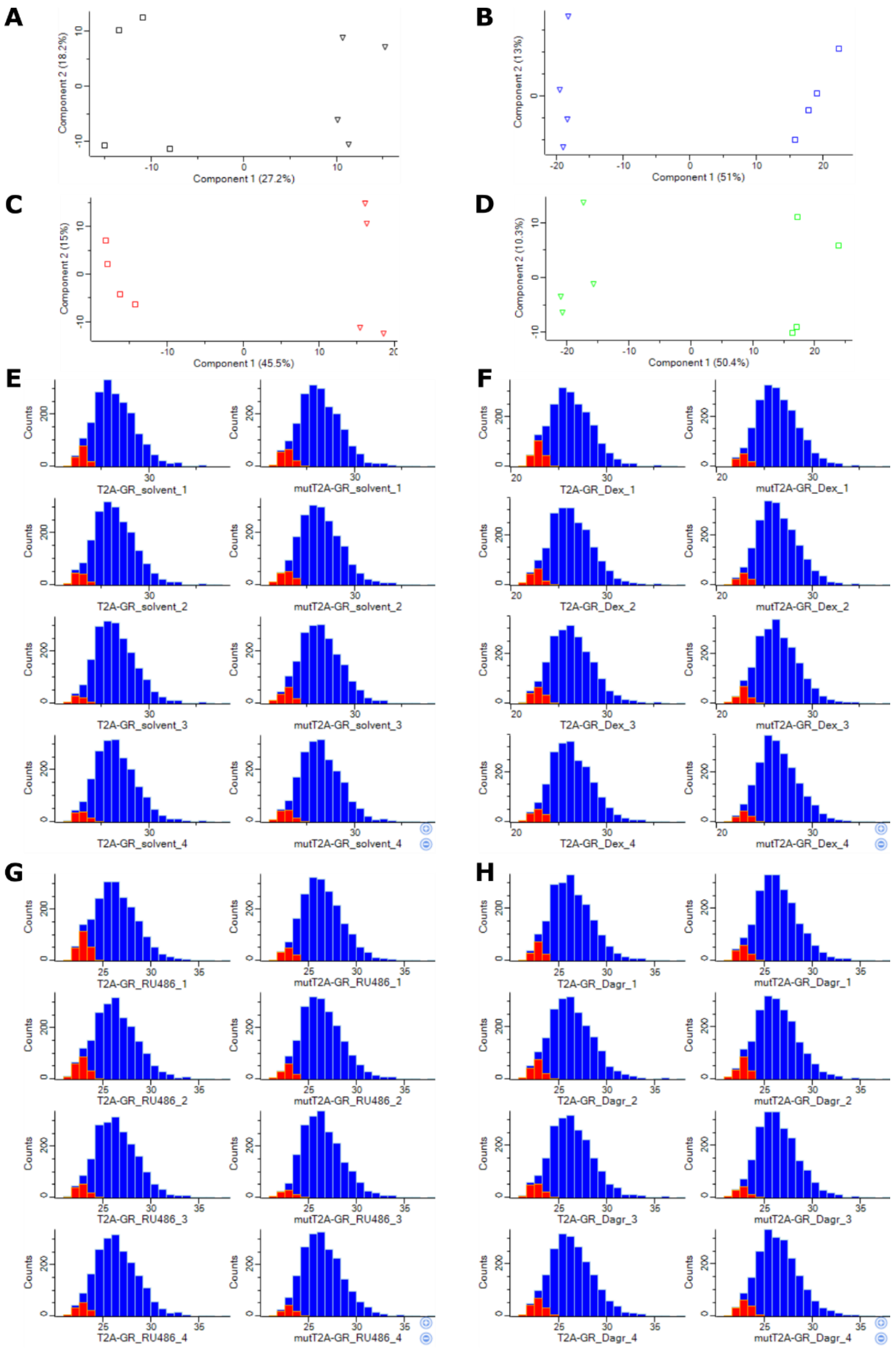


**Fig. S1**. Histograms and PCA plots of each experimental condition. **(A-D)** Principal Component Analysis for **(A)** solvent, **(B)** Dex, **(C)** RU486, **(D)** Dagr made in Perseus. Category enrichment was enabled. Cut-off was determined using the Benjamini-Hochberg method at a 5% FDR, without relative enrichment. Triangles represent control cell line (A549 TurboID-T2A-GR), squares represent the target cell line (A549 TurboID-mutT2A-GR). **(E-H)** Histograms representing the distribution of $log_2$(LFQ) values for **(E)** solvent, **(F)** Dex, **(G)** RU486 and **(H)** Dagr samples. Imputed values are indicated in red.

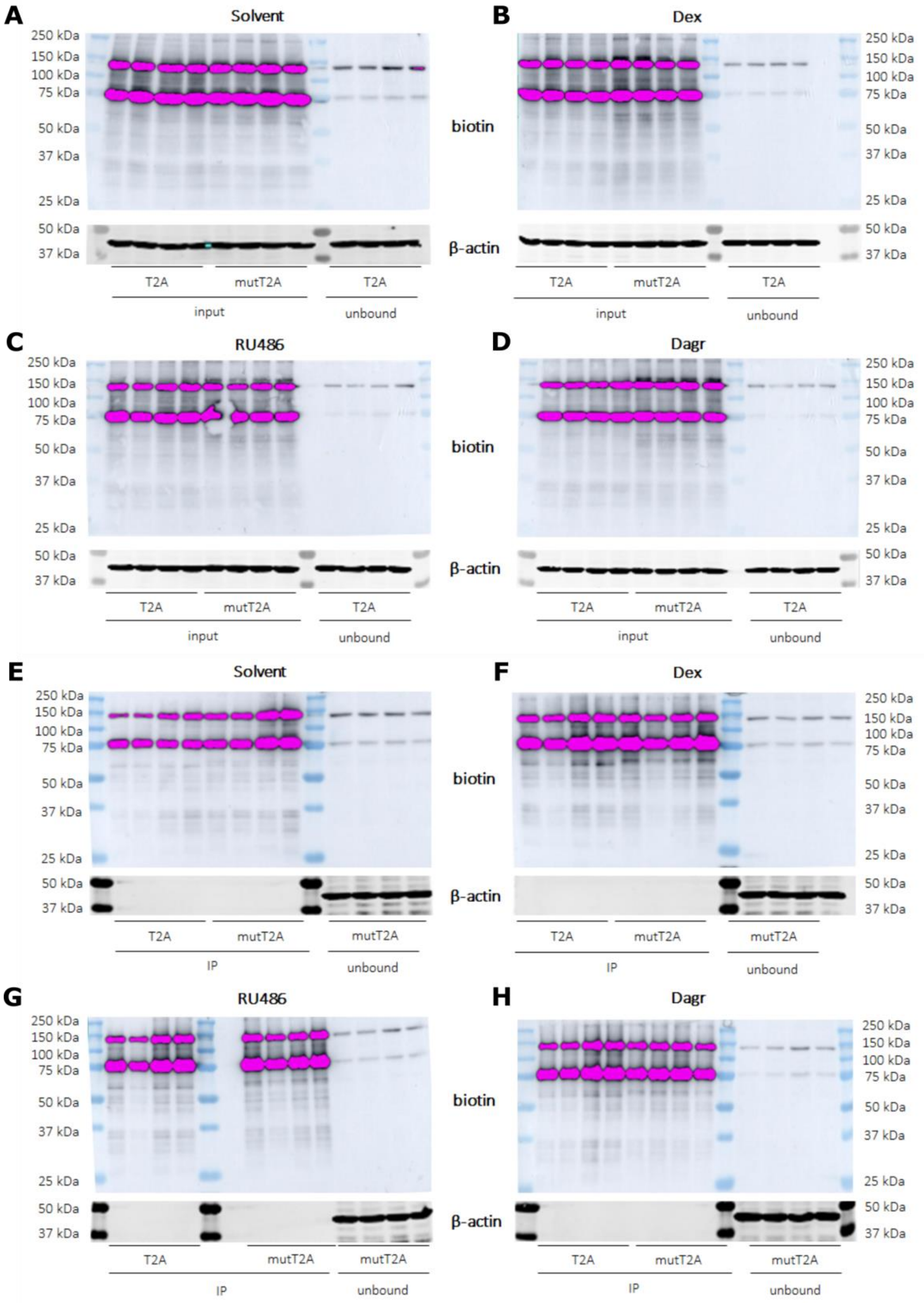


**Fig. S2.** Immunoblot analysis of input, immunoprecipitated and unbound fractions of all conditions following detection with anti-β-actin and streptavidin-HRP.

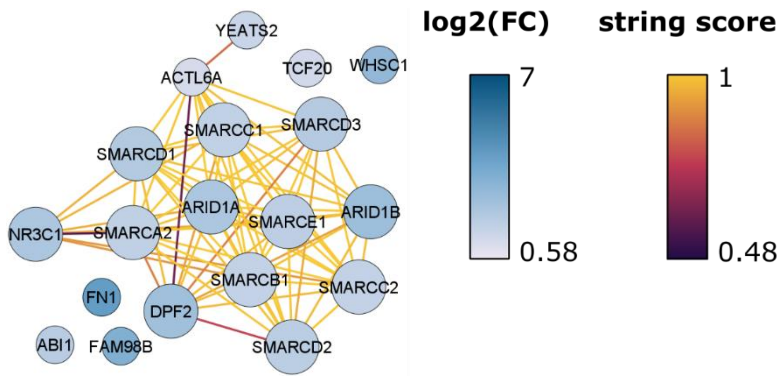


**Fig. S3**. Physical STRING network of the significantly enriched GR interactors following 2 h solvent treatment. Small circles represent significantly enriched proteins at a 5% FDR, larger circles represent significantly enriched proteins at a 1% FDR. The blue color scale indicates the $log_2$(fold change) between the target versus control cell line. The purple-yellow color scale corresponds to the confidence of the interaction according to the STRING database.

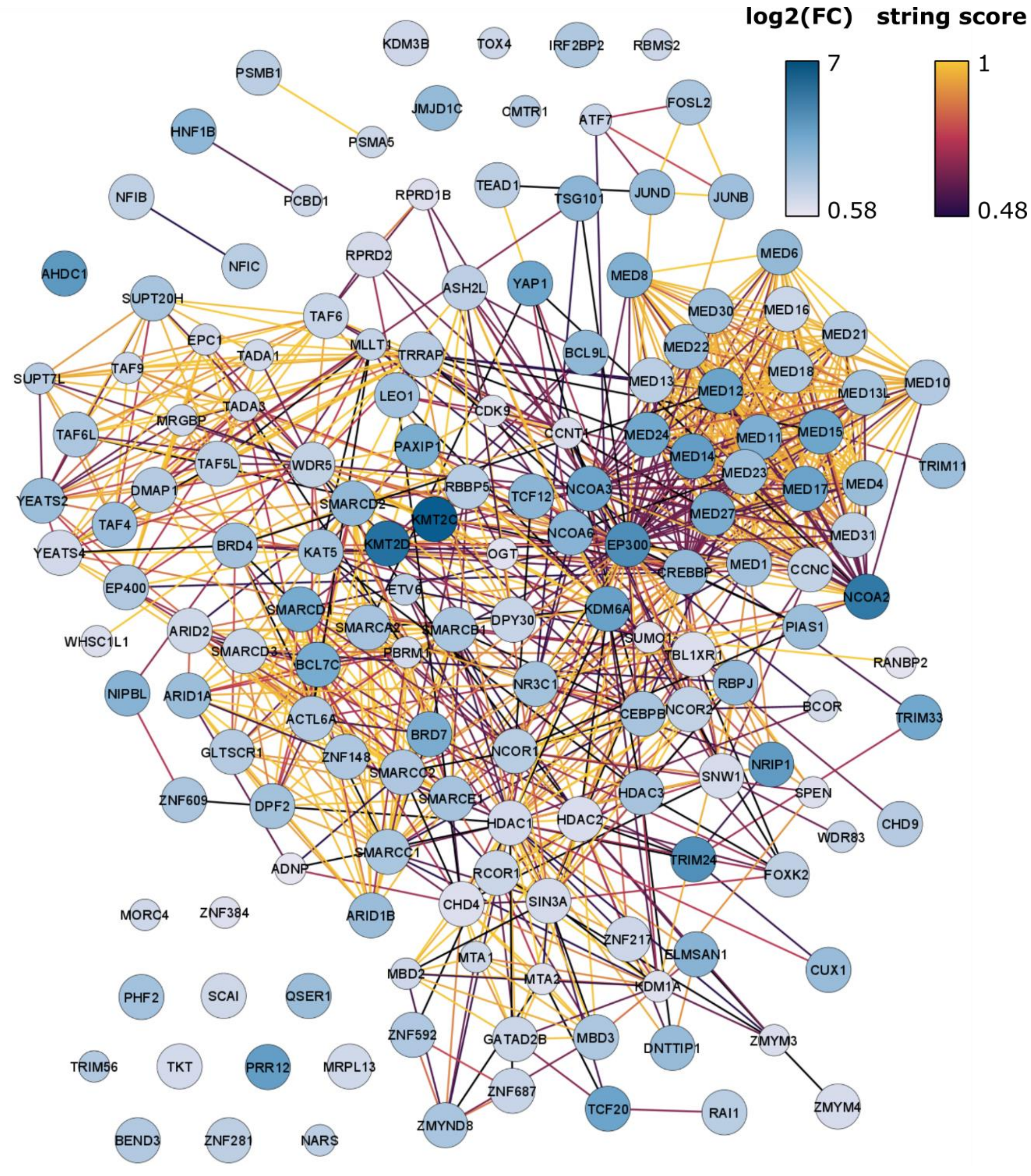


**Fig. S4**. Physical STRING network of the significantly enriched GR interactors following 2 h Dex treatment. Small circles represent significantly enriched proteins at a 5% FDR, larger circles represent significantly enriched proteins at a 1% FDR. The blue color scale indicates the $\log_2$(fold change) between the target versus control cell line. The purple-yellow color scale corresponds to the confidence of the interaction according to the STRING database.

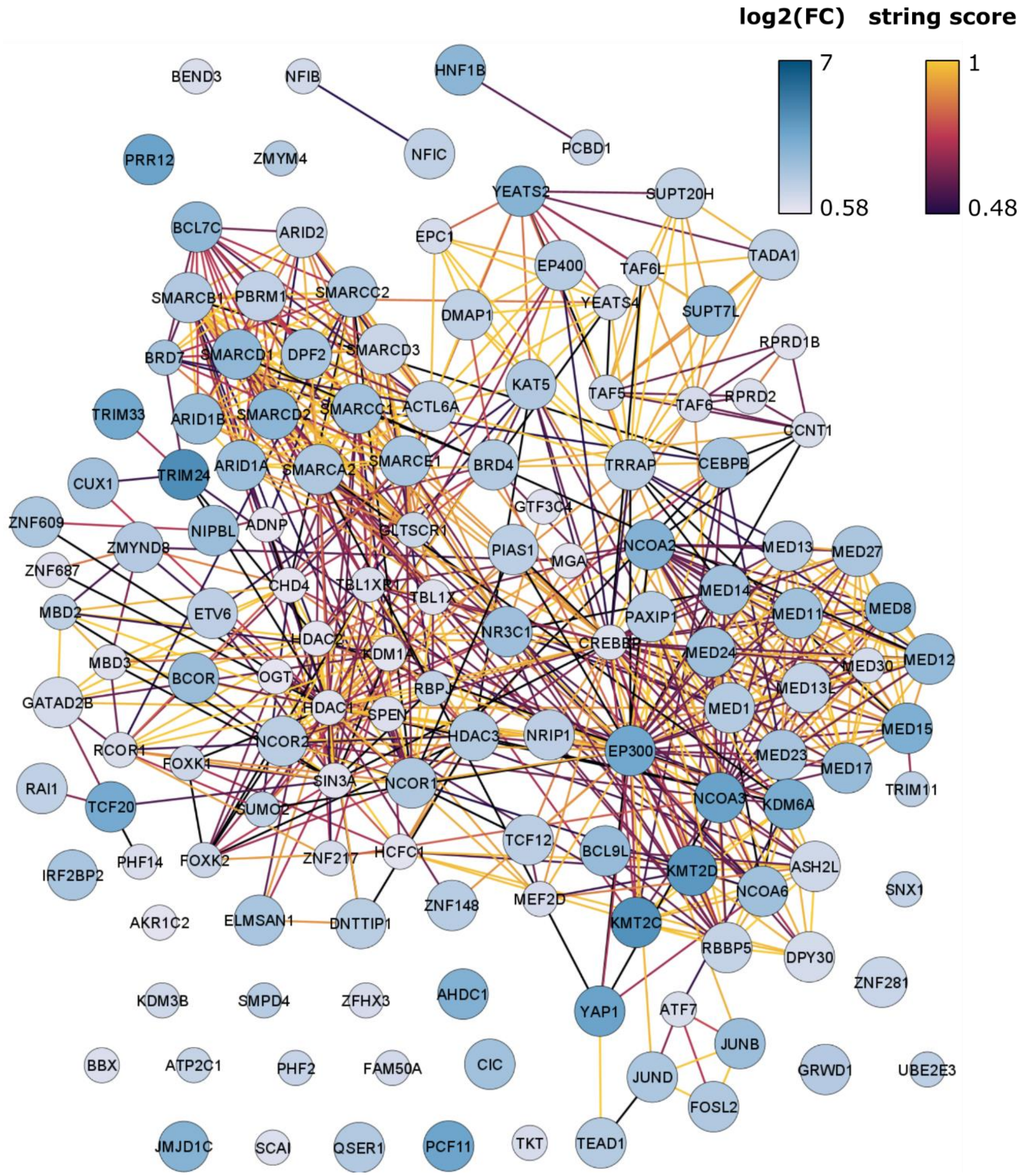


**Fig. S5**. Physical STRING network of the significantly enriched GR interactors following 2 h RU486 treatment. Small circles represent significantly enriched proteins at a 5% FDR, larger circles represent significantly enriched proteins at a 1% FDR. The blue color scale indicates the $\log_2$(fold change) between the target versus control cell line. The purple-yellow color scale corresponds to the confidence of the interaction according to the STRING database.

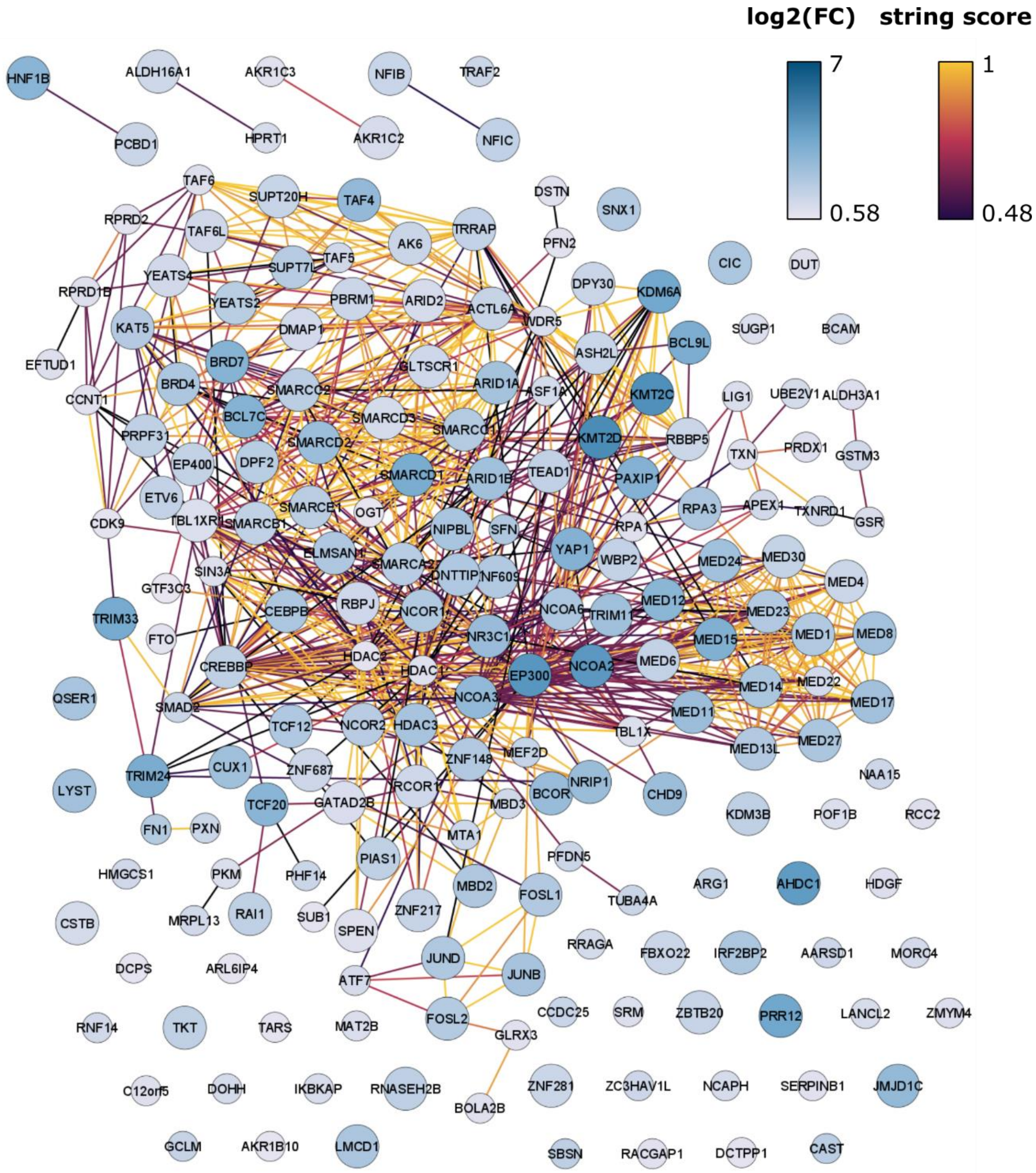


**Fig. S6**. Physical STRING network of the significantly enriched GR interactors following 2 h Dagr treatment. Small circles represent significantly enriched proteins at a 5% FDR, larger circles represent significantly enriched proteins at a 1% FDR. The blue color scale indicates the $log_2$(fold change) between the target versus control cell line. Purple-yellow color scale corresponds to the confidence of the interaction according to the STRING database.

**Table S2**. Overview of significantly enriched proteins across all conditions (FDR 0.01; $S_0$=0.58), ranked according to descending $log_2$(target/control) values in the Dex condition. Bait is indicated in bold italics, known GR interactors in BioGRID are indicated in bold. Da, Dagrocorat; Dex, Dexamethasone; RU, RU486; So, solvent.

| Gene name | So | Dex | RU | Da | Gene name | So | Dex | RU | Da | Gene name | So | Dex | RU | Da |
|---|---|---|---|---|---|---|---|---|---|---|---|---|---|---|
| KMT2C | | x | x | x | JUNB | | x | x | x | **NFIB** | | x | | x |
| **KMT2D** | | x | x | x | MED30 | | x | | x | **CIC** | | | x | x |
| **NCOA2** | | x | x | x | **MED1** | | x | x | x | CCNC | | x | | |
| **EP300** | | x | x | x | LEO1 | | x | | | **NCOR2** | | x | x | x |
| **TRIM24** | | x | x | x | **KAT5** | | x | x | x | WDR5 | | x | | |
| AHDC1 | | x | x | x | PHF2 | | x | | | **DPY30** | | x | x | x |
| **NRIP1** | | x | x | x | BRD4 | | x | x | x | **ZNF687** | | x | | x |
| **MED14** | | x | x | x | **DPF2** | x | x | x | x | TAF6 | | x | | |
| PRR12 | | x | x | x | ***NR3C1*** | x | x | x | x | TAF9 | | | | x |
| **KDM6A** | | x | x | x | **SMARCA2** | x | x | x | x | GATAD2B | | x | x | x |
| **MED17** | | x | x | x | **SMARCE1** | x | x | x | x | **RCOR1** | | x | | x |
| MED12 | | x | x | x | HDAC3 | | x | x | x | MED16 | | x | | |
| **NCOA3** | | x | x | x | MED13L | | x | x | x | PCBD1 | | | | x |
| **TCF20** | | x | x | x | **CEBPB** | | x | x | x | SMARCD3 | x | x | x | x |
| YAP1 | | x | x | x | SUPT20H | | x | x | x | **KDM3B** | | x | | x |
| **MED15** | | x | x | x | ZMYND8 | | x | x | | ZNF217 | | x | | x |
| TRIM33 | | x | x | x | ZNF609 | | x | x | x | MBD2 | | | | x |
| MED24 | | x | x | x | **SMARCC1** | x | x | x | x | SCAI | | x | | |
| **SMARCD1** | x | x | x | x | FOSL2 | | x | x | x | **ARID2** | | x | x | x |
| BCL7C | | x | x | x | TAF6L | | x | | x | MRPL13 | | x | | |
| **PAXIP1** | | x | x | x | **SMARCB1** | x | x | x | x | YEATS4 | | x | | x |
| BRD7 | | x | | x | MBD3 | | x | | | TADA1 | | | x | |
| MED27 | | x | x | x | ZNF148 | | x | x | x | TKT | | x | | x |
| MED11 | | x | x | x | **SMARCC2** | x | x | x | x | **PBRM1** | | | x | x |
| **MED8** | | x | x | x | **IRF2BP2** | | x | x | x | **HDAC1** | | x | | |
| **NIPBL** | | x | x | x | ZNF592 | | x | | | **BCOR** | | | x | x |
| ELMSAN1 | | x | x | x | EP400 | | x | x | x | SIN3A | | x | | |
| **NCOA6** | | x | x | x | MED18 | | x | | | RPRD2 | | x | | |
| **TSG101** | | x | | | BEND3 | | x | | | SNW1 | | x | | |
| TCF12 | | x | x | x | MED21 | | x | | | ZMYM4 | | x | | |
| HNF1B | | x | x | x | ACTL6A | | x | x | x | **HDAC2** | | x | | |
| BCL9L | | x | x | x | MED10 | | x | | | TBL1XR1 | | x | | x |
| SMARCD2 | x | x | x | x | CHD9 | | x | | x | CHD4 | | x | | |
| MED6 | | x | | x | DMAP1 | | x | x | x | **SPEN** | | | | x |
| **CREBBP** | | x | | x | GLTSCR1 | | x | | x | FBXO22 | | | x | |
| RBPJ | | x | | x | RBBP5 | | x | x | x | AKR1C2 | | | | x |
| **JMJD1C** | | x | x | x | **SUPT7L** | | | x | x | ALDH16A1 | | | | x |
| **MED22** | | x | | | NFIC | | x | x | x | PRPF31 | | | | x |
| DNTTIP1 | | x | x | x | **RAI1** | | x | x | x | WBP2 | | | | x |
| **CUX1** | | x | x | x | **NCOR1** | | x | x | x | LMCD1 | | | | x |
| JUND | | x | x | x | TRRAP | | x | x | x | SNX1 | | | | x |
| MED4 | | x | | x | TEAD1 | | x | x | x | CSTB | | | | x |
| QSER1 | | x | x | x | MED13 | | x | x | | RNASEH2B | | | | x |
| **ARID1B** | x | x | x | x | PSMB1 | | x | | | PCF11 | | | x | |
| TAF4 | | x | | x | ETV6 | | | x | x | FOSL1 | | | | x |
| **YEATS2** | | x | x | x | **ZNF281** | | x | x | x | GRWD1 | | | x | |
| **ARID1A** | | x | x | x | **ASH2L** | | x | x | x | LYST | | | | x |
| **PIAS1** | | x | x | x | TAF5L | | x | | | **ZBTB20** | | | | x |
| MED23 | | x | x | x | FOXK2 | | x | | | | | | | |
| TRIM11 | | x | | x | MED31 | | x | | | | | | | |

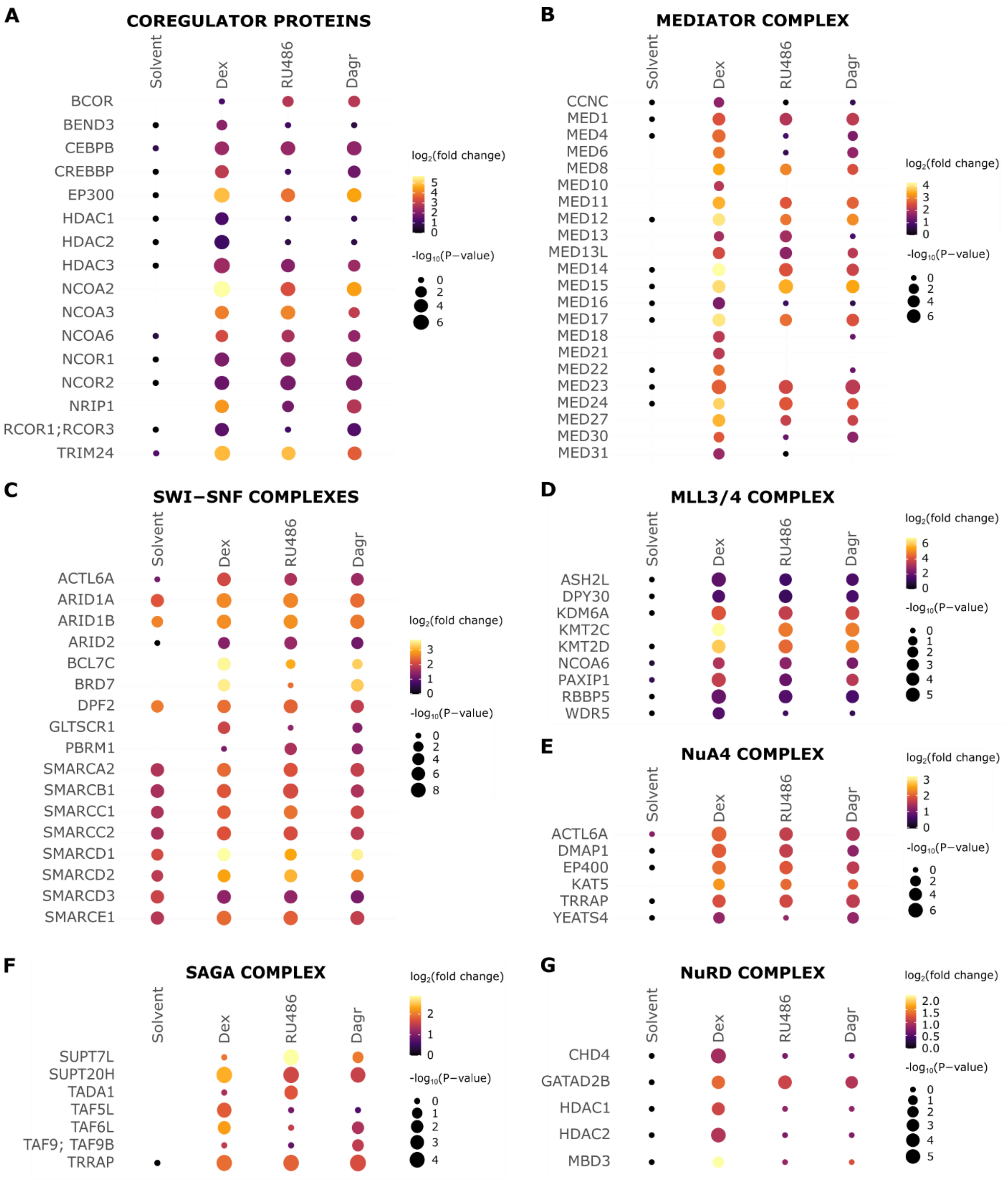


**Fig. S7**. Comparison of protein complex enrichments between all conditions. Dot plots represent relative enrichment of different protein complexes across conditions. Dot sizes correspond to -$\log_{10}$(P-values), with the smallest dot corresponding to proteins that were identified but not significantly enriched in target versus control cell lines. Color scale represents $\log_2$ fold-enrichments, with black dots representing proteins which were identified but less than 1.5-fold enriched.

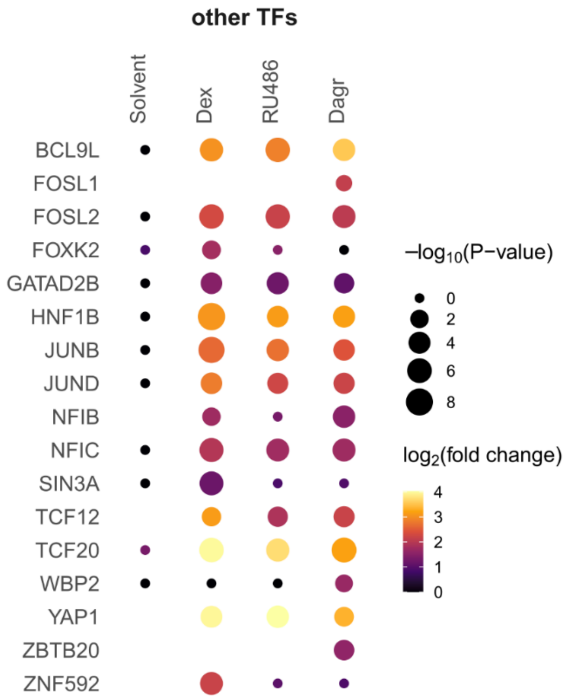


**Fig. S8**. Comparison of protein complex enrichments between all conditions. Dot plots representing relative enrichment of different protein complexes across conditions. Dot sizes correspond to $-\log_{10}$(P-values), with the smallest dot corresponding to proteins that were identified but not significantly enriched in target versus control cell lines. Color scale represents $\log_2$-fold enrichments, with black dots representing proteins that were identified but less than 1.5-fold enriched. TF, transcription factor.
